# Nectar-dwelling microbes of common tansy are attractive to its mosquito pollinator, *Culex pipiens* L

**DOI:** 10.1101/2020.04.03.024380

**Authors:** D. A. H. Peach, C. Carroll, S. Meraj, S. Gomes, E. Galloway, A. Balcita, H. Coatsworth, N. Young, Y. Uriel, R. Gries, C. Lowenberger, M. Moore, G. Gries

## Abstract

**Background:** There is widespread interkingdom signalling between insects and microbes. For example, microbes found in floral nectar may modify its nutritional composition and produce odorants that alter the floral odor bouquet which may attract insect pollinators. Mosquitoes consume nectar and can pollinate flowers. We identified microbes isolated from nectar of common tansy, *Tanacetum vulgare,* elucidated the microbial odorants, and tested their ability to attract the common house mosquito, *Culex pipiens.*

**Results:** We collected 18 microbial isolates from *T. vulgare* nectar, representing at least 12 different taxa which we identified with 16S or 26S rDNA sequencing as well as by biochemical and physiological tests. Three microorganisms *(Lachancea thermotolerans, Micrococcus lactis, Micrococcus luteus)* were grown on culture medium and tested in bioassays. Only the yeast *L. thermotolerans* grown on nectar, malt extract agar, or in synthetic nectar broth significantly attracted *C. pipiens* females. The odorant profile produced by *L. thermotolerans* varied with the nutritional composition of the culture medium. Surprisingly, all three microbes grown separately, but presented concurrently, attracted fewer *C. pipiens* females than *L. thermotolerans* by itself.

**Conclusions:** Floral nectar of *T. vulgare* contains various microbes whose odorants contribute to the odor profile of inflorescences. In addition, *L. thermotolerans* produced odorants that attract *Cx. pipiens* females. As the odor profile of *L. thermotolerans* varied with the composition of the culture medium, we hypothesize that microbe odorants inform nectar-foraging mosquitoes about the availability of certain macro-nutrients which, in turn, affect foraging decisions by mosquitoes.

## Introduction

Signalling between microbes and insects is widespread and occurs in a variety of contexts [1–3]. Plant odorants as well as visual displays of inflorescences play essential roles in attracting insect pollinators [4]. Floral nectar provides nutrition and habitat for myriad microorganisms [5–10] that may alter the composition of nectar [11] and produce odorants [8, 12, 13], thereby modifying the inflorescence odor bouquet [13–15]. These microbially-derived odorants may contribute to the plant-pollinator signalling system by serving as attractive semiochemicals (message-bearing chemicals) to pollinators [13, 16–18]. However, this type of signalling may be species-specific with respect to both the sender and the receiver of these semiochemicals because in other instances, microbe-derived odorants cause no behavioral response [19], or avoidance by pollinators [19, 20].

Plant-derived nutrients (e.g., sugars) are fundamental dietary constituents for adult mosquitoes [21], providing energy for flight, mating, blood-feeding, egg-laying, and female overwintering [21–23]. Floral nectar is the dominant source of plant sugar for most mosquitoes but other sugar sources such as extra-floral nectar, aphid honeydew, and fruit juices are also consumed [21, 24]. Inflorescence semiochemicals [25] along with visual inflorescence displays [26] and CO_2_ [27] attract mosquitoes to various inflorescences [21, 27–29] that they discern [30, 31] and may pollinate [32–35].

Microbe-derived odorants have not yet been implicated in mosquito nectar-foraging but are exploited by mosquitoes in a variety of other contexts. For example, the odor bouquet of microbe-inoculated or infested aphid honeydew is more attractive to the yellow fever mosquito, *Aedes aegypti,* than the odor bouquet of sterilized honeydew [36]. The volatile semiochemicals emitted by human skin microbes help attract host-seeking mosquitoes [37–39]. Carbon dioxide is another important vertebrate- and plant-host cue for mosquitoes [27, 40], which originates not only from potential hosts but also from their symbiotic microbes [41]. Moreover, microbe-derived semiochemicals indicate suitable oviposition sites for many mosquito species and attract gravid females [42–44]. Microbe-derived semiochemicals and CO_2_ may also play a role in mosquito attraction to floral nectar.

Working with the common tansy, *Tanacetum vulgare*, and one of its mosquito pollinators, the common house mosquito, *Culex pipiens* [35], we tested the hypotheses (H) that: (1) nectar-colonizing microbes emit semiochemicals attractive to *Cx. pipiens;* (2) the attractiveness of these microbes is dependent upon their nutrient source; and (3) multiple species of nectar-colonizing microbes attract more mosquitoes than a single species.

## Results

### Identification of Nectar-Colonizing Microbes

We collected nectar from nectaries of *T. vulgare* florets with a sterile glass microcapillary tube and identified 18 microbial isolates (Table 1) by sequencing the 16S or 26S rDNA genes, and by comparing the results to data in the National Center for Biotechnology Information GenBank using BLASTn (Bethesda, USA; http://www.ncbu.nlm.nih.gov/BLAST.cgi). We performed additional biochemical and physiological tests on select isolates to aid in their identification (Table 2). The yeast *L. thermotolerans* was present in nectar from two separate florets on two separate plants. *Bacillus* spp. were present in six florets from three separate plants, and *Micrococcus* spp. were present in four florets from three separate plants. In five cases, more than one microbe occurred in the same floret.

**Table 1.** (*i*) List of microbes identified from *Tanacetum vulgare* nectar including information about the plant, inflorescence, and individual florets from which they were collected, *(ii)* the medium used to culture microbes, and *(iii)* the methods used for microbe identification. LB = Luria-Bertani; YEPD = yeast extract peptone dextrose; ID = identification; PCR = polymerase chain reaction; test = traditional biochemical and physiological tests.

| Isolate | Plate Media | Name of Microbe | Source |  |  |  |
| --- | --- | --- | --- | --- | --- | --- |
|  |  |  | Plant # | Inflorescence # | Floret # | ID Method |
| LB-T1D1 | LB | Mix of *Bacillus amyloliquefaciens* & *B. subtilis* | 1 | 1 | 1 | PCR + test |
| LB-T1E1 | LB | *Pseudarthrobacter* sp. | 1 | 2 | 1 | PCR |
| LB-T1E2 | LB | *Micrococcus lactis* | 1 | 2 | 2 | PCR + test |
| LB-T2C1 | LB | *Pseudomonas rhizosphaerae* | 2 | 1 | 1 | PCR |
| LB-T2E1 | LB | *Staphylococcus epidermidis* | 2 | 1 | 1 | PCR |
| LB-T3A1 | LB | *Cryptococcus* sp. | 3 | 1 | 1 | PCR |
| LB-T3B2 | LB | *Bacillus* sp. | 3 | 2 | 1 | PCR |
| LB-T4A2 | LB | *Rhodotorula/Ustilentyloma* sp. | 4 | 1 | 1 | PCR |
| LB-T4B1 | LB | *Bacillus circulans* | 4 | 2 | 1 | PCR + test |
| LB-T4B2 | LB | *Bacillus* sp. | 4 | 2 | 2 | PCR |
| LB-T4C1 | LB | *Micrococcus luteus* | 4 | 3 | 1 | PCR+ test |
| LB-T4D1 | LB | *Bacillus* sp. | 4 | 3 | 1 | PCR |
| YPD-T1B1 | YEPD | *Lachancea thermotolerans* | 1 | 3 | 1 | PCR |
| YPD-T1E1 | YEPD | *Micrococcus* sp. | 1 | 2 | 1 | PCR |
| YPD-T2E1 | YEPD | *Lachancea thermotolerans* | 2 | 2 | 2 | PCR + test |
| YPD-T3A1 | YEPD | *Micrococcus* sp. | 3 | 1 | 1 | PCR |
| YPD-T3B2 | YEPD | *Bacillus* sp. | 3 | 2 | 1 | PCR |
| YPD-T4C1 | YEPD | *Bacillus* sp. | 4 | 3 | 1 | PCR |

**Table 2.** Biochemical tests for the identification of microbes collected from the inflorescences of common tansy, *Tanacetum vulgare.*

| Test | YPD-T2E1 | LB-T1D1-a<br>(round) | LB-T1D1-b<br>(filamentous) | LB-T4B1 | LB-T1E2 | LB-T4C1 |
| --- | --- | --- | --- | --- | --- | --- |
| *Fermentation of:* |  |  |  |  |  |  |
| Sucrose | + | + | + | + | Not tested | Not tested |
| Starch | + | + | + | + | Not tested | Not tested |
| L-alanine | - | - | - | - | Not tested | Not tested |
| Glucose | + | + | + | + | Not tested | Not tested |
| Mannitol | - | + | + | + | Not tested | Not tested |
| Xylose | - | - | - | - | Not tested | Not tested |
| Galactose | + | - | - | + | Not tested | Not tested |
| Lactose | - | + | + | Varies | Not tested | Not tested |
| Fructose | + | + | + | + | Not tested | Not tested |
| *Enzyme production:* |  |  |  |  |  |  |
| Catalase test | + | + | + | + | Not tested | Not tested |
| Oxidase test | Not tested | - | - | Varies | Not tested | Not tested |
| Other tests: |  |  |  |  | Not tested | Not tested |
| 7% NaCl development | Not tested | + | + | - | - | Not tested |
| 10% NaCl development | Not tested | + | + | - | Not tested | Not tested |
| Nitrate reduction | Not tested | Not tested | Not tested | Not tested | + | + |
| Hydrolysis of casein | Not tested | Not tested | Not tested | Not tested | + | Not tested |
| Hydrolysis of tween80 | Not tested | Not tested | Not tested | Not tested | Not tested | - |
| Assimilation of L-arabinose | Not tested | Not tested | Not tested | Not tested | - | Not tested |
| Assimilation of D-glucose | Not tested | Not tested | Not tested | Not tested | - | Not tested |
| 10 °C development | Not tested | Not tested | Not tested | Not tested | + | - |
| 42 °C development | Not tested | Not tested | Not tested | Not tested | - | Not tested |
| 50 °C development | Not tested | - | + | - | Not tested | Not tested |
| KOH string test | Not relevant | Gram + | Gram + | Gram + | Not tested | Not tested |

### H1: Nectar-colonizing microbes emit semiochemicals attractive to Cx pipiens

In two-choice laboratory experiments with a paired-trap design, we tested attraction of female *Cx. pipiens* to a synthetic nectar broth (10% w/v sucrose, 2% w/v yeast extract) (control stimulus) and the same broth inoculated with (*i*) *L. thermotolerans* (Exp. 1), (*ii*) *M. luteus* (Exp. 2), or (*iii*) *M. lactis* (Exp. 3). We captured more female *Cx. pipiens* when *L. thermotolerans* was the inoculum (z = 4.03, p < 0.0001; Fig. 1, Exp. 1) but not when either *M. luteus* or *M. lactis* was the inoculum (*M. luteus:* z = 1.44, p = 0.15; *M. lactis:* z = −1.02, p = 0.31; Fig 1, Exps. 2, 3), indicating an ability of the mosquitoes to discern among different microbes or their metabolites.

**Fig 1.**
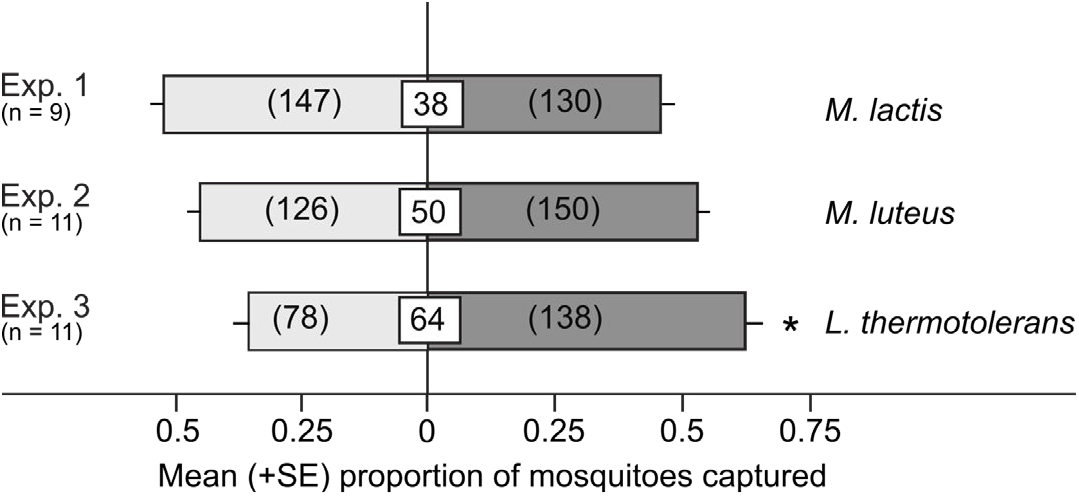
Mean proportion of *Culex pipiens* females captured in paired traps baited with a synthetic nectar broth inoculated, or not (control; light grey bars), with *Micrococcus lactis* (Exp. 1), *Micrococcus luteus* (Exp. 2), or *Lachancea thermotolerans* (Exp. 3). Numbers in white boxes represent the mean percentage of non-responding mosquitoes, and numbers within parentheses the total number of mosquitoes captured. The asterisk (*) in Exp. 3 indicates a significant preference for the treatment stimulus (binary logistic regression with a logit link function, P < 0.05).

### Dose of Microbes Tested

We grew *L. thermotolerans*, *M. luteus* and *M. lactis* in the synthetic nectar broth for 48 h and these reached final mean concentrations of 3.85 × 10^7^ cells/mL (N = 2), 1.13 × 10^5^ cells/mL (N = 2), and 2.85 × 10^6^ cells/mL (N = 2), respectively.

### H2: The attractiveness of microbes is dependent upon their growth medium

To determine whether the attractiveness of *L. thermotolerans* is affected by its nutrient source, single colonies of *L. thermotolerans* were spread-plated onto synthetic nectar agar, malt extract agar or YEPD agar plates. In two-choice laboratory experiments, we then tested attraction of *Cx. pipiens* to paired traps baited with either one of the three types of inoculated agar or an uninoculated control agar. *Lachancea thermotolerans* growing on synthetic nectar agar (Exp. 4) or malt extract agar (Exp. 5) attracted more female *Cx. pipiens* than corresponding agar controls but not when growing on YEPD agar (Exp. 6) (Exp. 4: z = 2.29, p = 0.02; Exp. 5: z = 2.47, p = 0.013; Exp. 6: z = – 0.61, p = 0.54; Fig. 2). Thus, attraction of *Cx. pipiens* females to *L. thermotolerans* is contingent upon the nutrients available to this yeast.

**Fig 2.**
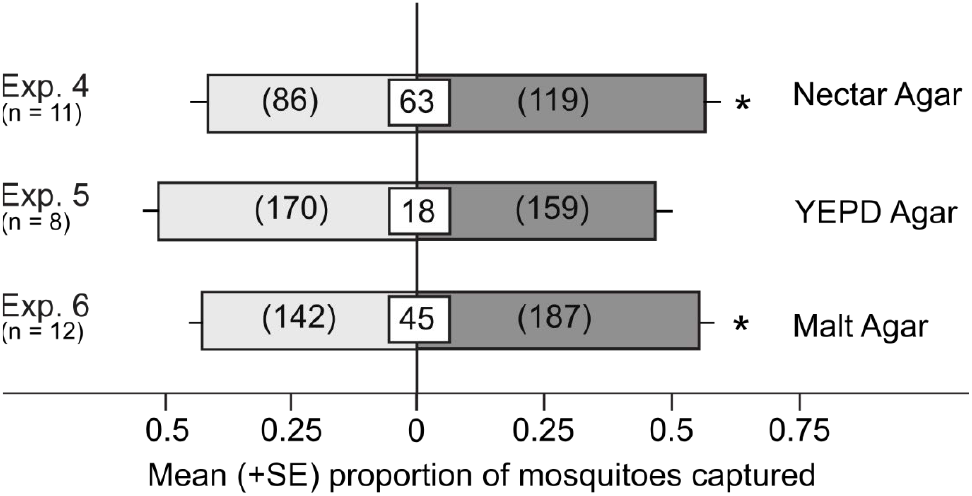
Mean proportion of *Culex pipiens* females captured in paired traps baited with one of three types of media inoculated, or not (control; light grey bars), with *Lachancea thermotolerans.* Numbers in white boxes represent the mean percentage of non-responding mosquitoes, and numbers within the parentheses the total number of mosquitoes captured. The asterisk (*) in Exp. 4 and Exp. 6 indicates a significant preference for the treatment stimulus (binary logistic regression with a logit link function, P < 0.05).

### H3: Multiple species of nectar-colonizing microbes attract more mosquitoes than single microbe species

We hypothesized that odorants from different microbes may have additive or synergistic effects on attraction of *Cx. pipiens* females; therefore, we investigated whether *M. lactis, M. luteus* and *L. thermotolerans* presented together are more attractive than each microbe on its own. We inoculated synthetic nectar in separate petri dishes with single colonies of *M. lactis, M. luteus or L. thermotolerans,* and in laboratory experiments tested attraction of female *Cx. pipiens* to paired traps baited with each species alone or in ternary combination. Surprisingly, the ternary combination was as attractive as *M. lactis* alone (z = −1.08, p = 0.28; Fig. 3, Exp. 7) and *M. luteus* alone (z = −0.33, p = 0.74; Fig. 3, Exp. 8), and even less attractive than *L. thermotolerans* alone (z = 1.96, p = 0.05; Fig. 3, Exp. 9). Hence, the attractiveness of *L. thermotolerans* was actually reduced when it was presented alongside the two bacterial species.

**Fig 3.**
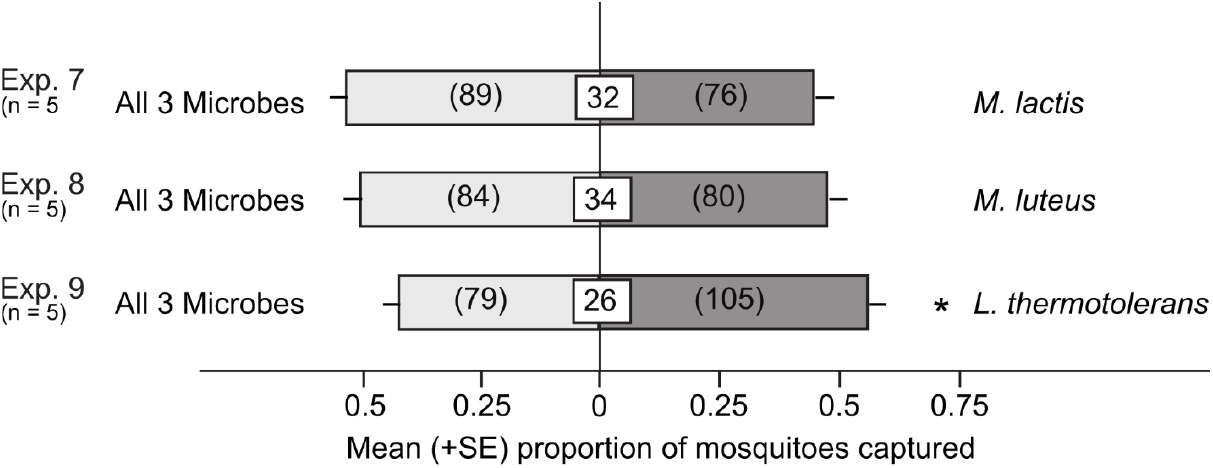
Mean proportion of *Culex pipiens* females captured in paired traps baited with one or three vessels containing synthetic nectar broth, each inoculated with one of three microbes: *Micrococcus lactis, M. luteus* and *Lachancea thermotolerans*. Numbers in white boxes represent the mean percentage of non-responding mosquitoes, and numbers within parentheses the total number of mosquitoes captured. The asterisk (*) in Exp. 9 indicates a significant preference for the *L. thermotolerans* test stimulus (binary logistic regression with a logit link function, P < 0.05).

### Identification of microbe-derived volatiles

We used dynamic headspace aerations to capture the odorants emitted from *L. thermotolerans* and identified them by gas chromatography-mass spectrometry (GC-MS). In response to the nutrients provided by the three types of media, *L. thermotolerans* produced different odor blends (Fig. 4). The yeast grown on all three media generated 2-phenylethanol, and dimethyl trisulfide was detected in two media types. All microbe-produced compounds differed from those originating from the media themselves (Table 3).

**Fig 4.**
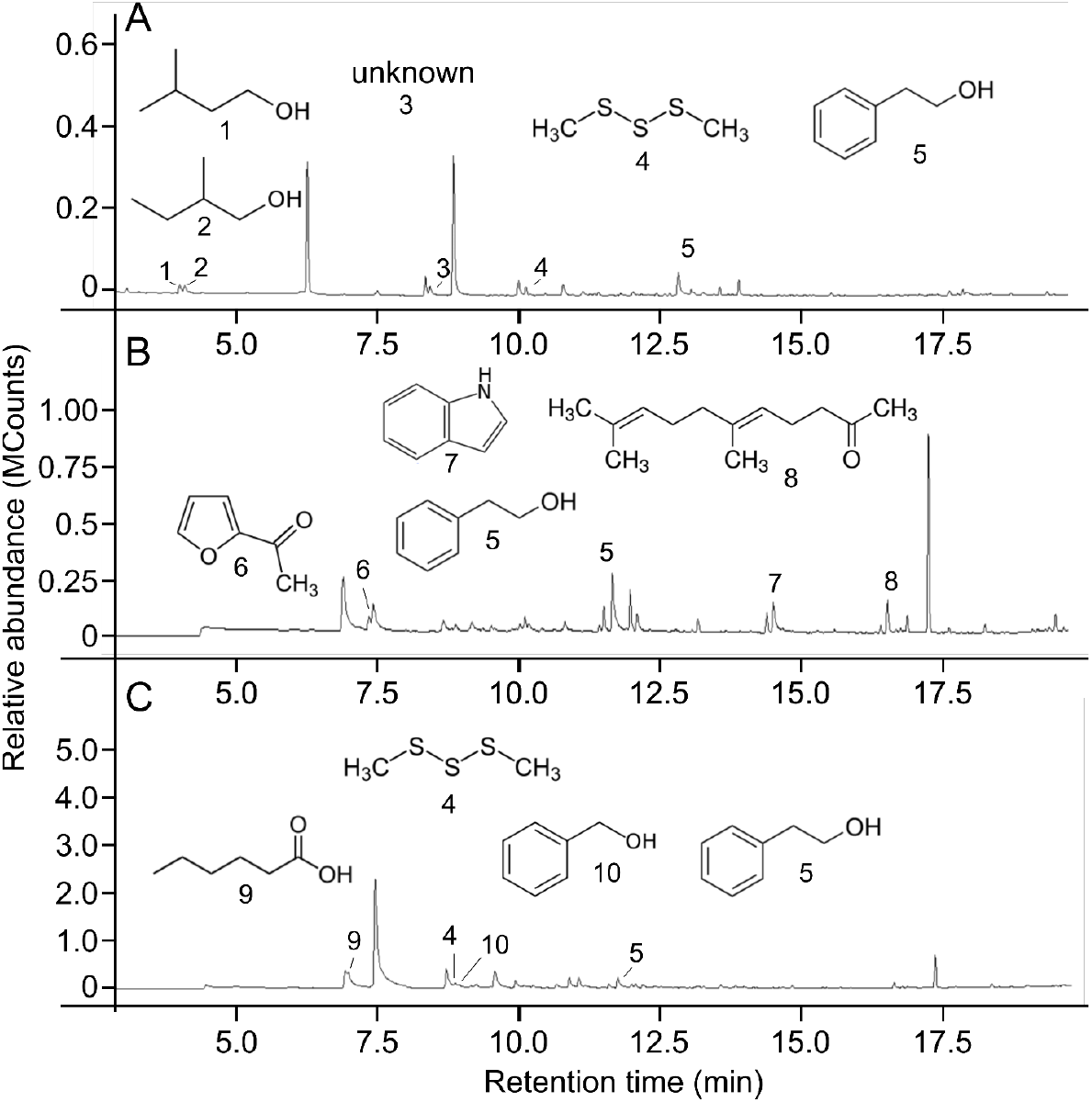
Headspace odorants from plates of YEPD agar (**A**), malt extract agar (**B**), and synthetic nectar agar plates (**C**) that were inoculated with *Lachancea thermotolerans* and that were not present in the headspace of uninoculated control plates. Compounds produced by *L. thermotolerans* were 3-methyl-butanol (1); 2-methyl-butanol (2); unknown (3); dimethyl trisulfide (4); 2-phenylethanol (5); 2-acetyl furan (6); indole (7); geranyl acetone (8); hexanoic acid (9) and benzyl alcohol (10). Note: different retention times of the same compounds in panels AC are due to different temperature programs run during gas chromatographic analyses (see Methods for details).

**Table. 3.** Headspace odorants of YEPD agar, malt extract agar and synthetic nectar agar, each inoculated with *Lachancea thermotolerans*. Compounds in bold font were found in two or more samples. ^a^All compounds were absent from the headspace of corresponding uninoculated control agar

| Medium | Compounds<sup>a</sup> |
| --- | --- |
| YEPD Agar | 2-Methyl-1-butanol<br>3-Methyl-1-butanol<br>**2-Phenylethanol**<br>**Dimethyl trisulfide** |
| Malt Extract Agar | 2-Acetyl furan<br>**2-Phenylethanol**<br>Indole<br>Geranyl acetone |
| Synthetic Nectar Agar | Hexanoic acid<br>**Dimethyl trisulfide**<br>Benzyl alcohol<br>**2-Phenylethanol** |

### CO_2_ production from synthetic nectar

We investigated the ability of *L. thermotolerans* to produce CO_2_ while growing in synthetic nectar broth sealed with a 98% sulfuric acid vapour lock. Over the course of 150 h, *L. thermotolerans* produced 343 mg of CO_2_ (Fig. 5).

**Fig. 5.**
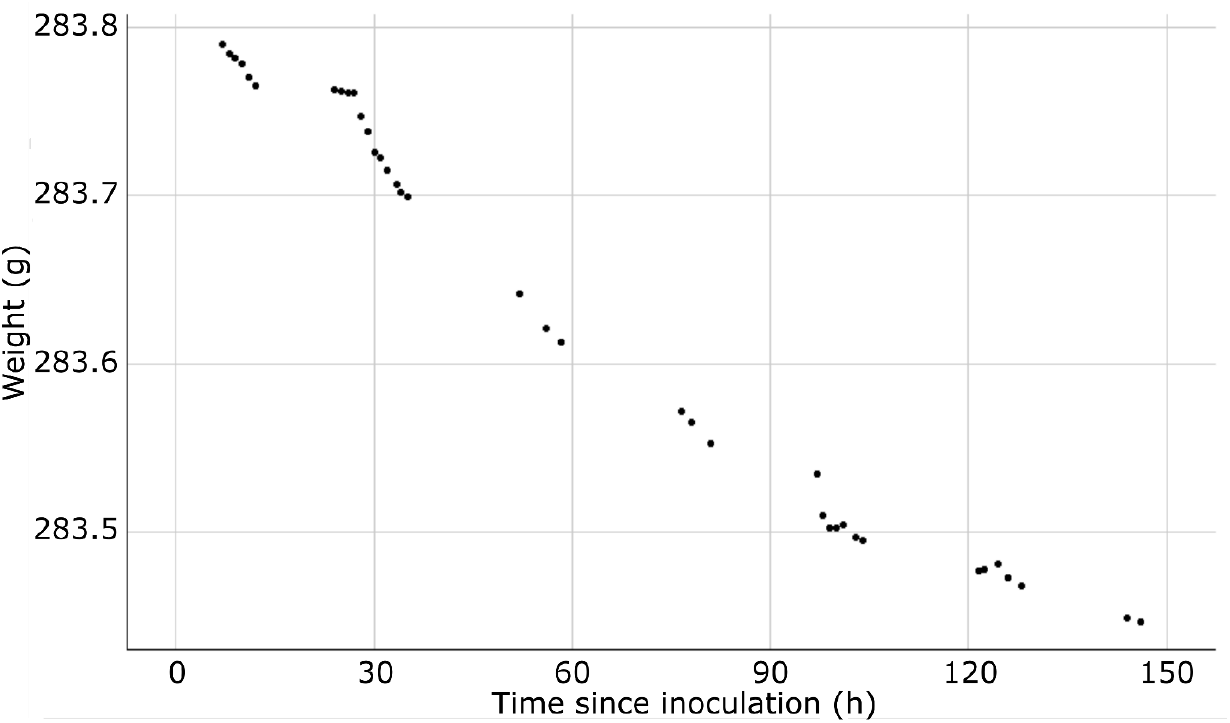
Weight loss over time of synthetic nectar broth (25 mL) inoculated with *Lachancea thermotolerans* due to fermentation and CO_2_ emission by *L. thermotolerans*.

## Discussion

Our data show that diverse microbes including the yeast *Lachancea thermotolerans* colonize floral nectar of tansy. *Lachancea thermotolerans* growing on synthetic nectar or malt extract media produce semiochemicals that attract female *Culex pipiens*. Furthermore, we show that *L. thermotolerans* grown in a synthetic nectar broth produces CO_2_ and that it is more attractive to female *Cx. pipiens* alone than when presented along with two bacteria also isolated from the same inflorescences. Below we elaborate on our conclusions.

Our culture-based approach to isolate nectar-colonizing microbes from common tansy likely under-represents the microbial diversity present in nectaries [45]; nevertheless, our collection of 18 isolates representing at least 12 species of microbes compares favourably with results of other studies examining microbial diversity in floral nectar using culture methods [6–8]. We isolated *Staphylococcus epidermidis* which may have been a contaminant because this microbe is typically associated with human skin. However, other microbes, such as *Candida albicans* [46] (not isolated in our study) which are thought to be obligate commensals of animals, have also been isolated from the environment [47]. Although culture-independent methods such as metagenomic analysis can be used to examine microbial diversity without the selection pressures of culturing, only by obtaining live cultures of the colonizing microbes could we study the odor profiles of these isolates and test their ability to attract *Cx. pipiens.*

Microbe-derived semiochemicals have been shown to guide foraging behavior of mosquitoes in a variety of contexts. Semiochemicals emitted from human skin microbiota, including *S. epidermidis*, *Corynebacterium minutissimum* and *Brevibacterium epidermidis*, attract mosquitoes to human hosts [37, 39, 48, 49]. Moreover, semiochemicals from *Psychrobacter immobilis*, *Sphingobacterium multivorum*, *Bacillus* spp., *Pseudomonas* spp., *Klebsiella* spp., and others help mosquitoes locate suitable oviposition sites [42, 44]. Finally, semiochemicals emitted from microbes colonizing aphid honeydew [36], or present in floral nectar [this study], attract sugar-foraging mosquitoes.

Microbes could also inform mosquitoes about the prospect of obtaining a sugar meal. The odor profiles of inflorescences differ not only between plant species [50] but also within the same species due, in part, to microbe-specific semiochemicals [17]. These semiochemicals may enable mosquitoes to discern nectar-rich and -poor inflorescences analogously to mosquitoes selecting human hosts based, in part, on their skin microbiota [38, 39]. Emission of microbe-semiochemicals from nectaries could inform mosquitoes about the presence of nectar-dwelling microbes which, in turn, would signal sugar or amino acid metabolism and thus the availability of sugar or amino acids. Alternatively, microbe semiochemicals could indicate microbe phoresis caused by a previous insect floral visit that may have temporarily depleted the sugar resource. Mosquitoes themselves are capable of microbe phoresis, as shown with floral nectar surrogates [51], as are many other insects visiting inflorescences or obtaining floral nectar [52, 53].

The informative value of microbe-derived odorants to foraging mosquitoes became evident when we grew *L. thermotolerans* in/on different nutrient sources. The nutrients available to *L. thermotolerans* not only affected the odorants it produced (Figs. 4–6) but also their attractiveness to foraging mosquitoes. This implies that the presence and composition of specific microbial odorants could inform mosquitoes about the availability of particular nutrients such as carbohydrates and amino acids. *Lachancea thermotolerans* grown in synthetic nectar broth produced appreciable amounts of CO_2_, which plays a significant role during both nectar- and host-foraging by mosquitoes [36].

*Lachancea thermotolerans* and its semiochemicals, respectively, also attract several species of North American yellowjackets (Hymenoptera: Vespidae) [54, 55] and the green lacewing, *Chrysoperla comanche* (Stephens) [56]. When grown and aerated on grape juice agar, *L. thermotolerans* produced 20 odorants which, when field-tested as a synthetic blend, attracted Western yellowjackets, *Vespula pensylvanica* [54]. Two of the odorants in this blend, 2-phentylethanol and 2-acetylfuran, were also found in this study. *Lachancea thermotolerans* is frequently isolated from fruit or fruit-related resources [57, 58] and is used in wine fermentations to generate ethanol and lactic acid [59].

Most studies investigating relationships between nectar-dwelling microbes and floral visitation by insects have focussed on hymenopterans (but see [60]), yet dipterans are also frequent visitors and important pollinators of flowers [61–64], and they interact with microbes [3, 65]. Our results suggest microbe-mediated, or at least modulated, inflorescence visitation by mosquitoes. This concept has been suggested for some hymenopteran pollinators including the European honey bee, *Apis mellifera* [13], and several species of bumble bees, *Bombus* spp. [17, 18], that preferentially visit inflorescences with nectar-dwelling yeasts, primarily *Metschnikowia reukaufii*. Conversely, both honey bees and bumble bees avoid inflorescences with certain nectar-dwelling bacteria [19, 20]. The combined information indicates that microbes can alter the floral scent [15], thereby prompting attraction or avoidance of specific floral visitors.

## Conclusion

We demonstrate that floral nectar of common tansies contains various microbes, and we identified 18 to genus level or further. Of the three species tested in behavioral bioassays, only the yeast *L. thermotolerans* had a significant effect on the attraction of female *Cx. pipiens,* which was diminished, rather than improved, by admixture with two bacterial species. The attractiveness of *L. thermotolerans* to *Cx. pipiens* females was dependent upon its nutrient source and linked to a distinct odorant profile, although a causal relationship was not tested. We propose that specific components of the odor blend signal the availability of certain macro-nutrients such as sugar and amino acids which, in turn, inform foraging decisions by mosquitoes.

## Materials and Methods

### Microbe collection

We collected 40 *T. vulgare* florets, from 20 inflorescences (2 florets per inflorescence), from 4 plants (5 inflorescences per plant) in Delta, BC (Canada), wearing latex gloves (VWR International, Radnor, USA) and a surgical mask (Acklands Ltd., Wawa, Canada). All plants were collected within a 3-m^2^ area beside a secondary road in a rural farming area. We immediately placed inflorescences into sterile Ziploc bags (S.C. Johnson and Son, Racine, USA) and stored them on ice for transport to the lab. Using ethanol and flame-sterilized scissors, we removed the tops of florets and inserted an autoclaved glass micro-capillary, prepared with a micropipette puller (Model P-1000, Sutter Instrument Co., Novato, USA), into the nectary to draw nectar via capillary action. Each draw yielded a maximum of ~1 μL of nectar. We repeated this twice on the same composite flower, using only one composite flower per inflorescence. We then inserted, and subsequently shattered, the micro-capillary into a sterile microcentrifuge tube (1.5 mL; ThermoFisher Scientific Inc., Waltham, USA) containing autoclaved distilled water (400 μL). We pipetted a 100-μL aliquot of this solution onto yeast extract peptone dextrose agar (YEPD) plates [66] and Luria-Bertani agar (LB) plates [67], spread it with a sterile glass rod, and then incubated plates for 48-72 h at 30 °C. We subsequently re-streaked morphologically distinct colonies onto new plates to obtain pure colonies. Working cultures were maintained at 4 °C. Storage cultures of each isolate were prepared with 20% glycerol and stored at −80 °C.

### Microbe identification

Cells from a single colony were picked and grown in liquid YEPD or LB media for 24 h at 30 °C, and DNA was extracted according to Rose et al. 1990 [68]. DNA concentrations were estimated using a NanoDrop UV/Vis 2000 spectrophotometer (ThermoFisher Scientific Inc., Waltham, USA). We used Taq DNA polymerase (Applied Biological Materials, Richmond, Canada) to amplify the V3-V4 loop of the 16S rDNA gene with the Universal Forward Primer (UniF) – 5’-CCTACGGGRBGCASCAG-3’ and the Universal Reverse Primer (UniR) – 5’-GGACTACNNGGGTATCTAAT-3’ [69], and to amplify the 26S rDNA gene with the NL1 primer – 5’-GCATATCAATAAGCGGAGGAAAAG-3’ – and NL4 primer – 5’-GGTCCGTGTTTCAAGACGG-3’ [70]. The identity of the colony found to be *L. thermotolerans* was confirmed using the specific primers INT2F (5’-TGGTTTTATTGAAGCCAAAGG-3’) and INT2R (5’-GGGGACCCGGAGATTAATAG-3’) [71]. PCR amplicons were pooled and concentrated using the NucleoSpin Gel and PCR Clean-up kit (Machery-Nagel, Duren, Germany). We sequenced amplicons (Genewiz, South Plainfield, USA) and used the Basic Local Alignment Search Tool (BLAST) [72] to compare the sequenced region of individual isolates with known sequences. A species or genus was determined to be a match if there was at least 95% coverage and 99% identity using BLAST with the sequenced isolate. In addition, we ran biochemical and physiological tests on select isolates (see Table 1) and Gram-tested bacterial colonies on select plates using KOH (Table 2). Biochemical tests were carried out according to established procedures [73, 74]. We determined the ability of isolates to grow at 7% NaCl and 10% NaCl in liquid media, and we tested isolate growth at various temperatures. For unknown *Bacillus* and *Epidermidis* isolates, catalase and oxidase enzyme production tests as well as sucrose, starch, L-alanine, glucose, mannitol, galactose, lactose, and fructose fermentation tests were run. For *Micrococcus* isolates, nitrate reduction, hydrolysis, and assimilation tests were run. We identified test isolates by comparing biochemical test results with data from the bioMérieux api^®^ 50 CHB/E test kit (bioMérieux SA, Lyon, France) and reference papers [75–79].

### Rearing of experimental mosquitoes

We reared *Cx. pipiens* at 23-26 °C, 40-60% RH, and a photoperiod of 14L:10D. We kept mixed groups of males and females in mesh cages (30 × 30 × 46 cm high) and provisioned them with a 10-% sucrose solution *ad libitum*. The primary author (DP) blood-fed females once per week. For oviposition, gravid females were given access to water in circular glass dishes (10 cm diameter × 5 cm high). We transferred eggs to water-filled trays (45 × 25 × 7 cm high) and sustained larvae with NutriFin Basix tropical fish food (Rolf C. Hagen Inc., Baie-D’Urfe, Canada). We transferred pupae via a 7-mL plastic pipette (VWR International, Radnor, USA) to water-containing 354-mL Solo cups covered with a mesh lid. Using an aspirator, we collected emergent adults and placed them in similar cups, along with a cotton ball soaked in a 10-% sucrose solution.

### Behavioural bioassays

We ran all behavioral bioassays in mesh cages (77 × 78 × 104 cm) which were wrapped in black fabric except for the top, thereby allowing ambient fluorescent light illumination. During bioassays, we kept cages at 23-26 °C, 40-60% RH and a photoperiod of 14L:10D. For each 24-h bioassay, we released 50 virgin *Cx. pipiens* females, 1-to 3-day-old, 24-h sugar-deprived into a cage fitted with adhesive-coated (The Tanglefoot Comp., Grand Rapids, USA) paired delta traps (9 cm × 15 cm) on burette stands spaced 30 cm apart. For each bioassay which ran for about 24 h, treatment and control stimuli were randomly assigned to these traps. At the end of each bioassay the number of mosquitoes in each trap was counted.

### Growth conditions for nectar-derived microorganisms

We grew microbes on YEPD agar plates, malt extract agar plates (3% w/v malt extract, 0.2% w/v peptone, 1.5% w/v agar), and synthetic nectar agar or nectar broth (10% w/v sucrose, 2% w/v yeast extract). All agar plates were 92-mm diam Petri dishes (Sarstedt Inc., Nümbrecht, Germany). The synthetic nectar broth was presented in an autoclaved 2-L Erlenmeyer flask. After streaking single colonies onto plates or inoculating broth, we incubated cultures for approximately 48-72 h at 23-26 °C and 40-60% RH. We used plates with microbial growth covering 40-60% of the surface area for behavioural bioassays.

### Attractiveness of microbes in synthetic nectar

In two-choice laboratory experiments with a paired-trap design, we tested attraction of female *Cx. pipiens* to microbe-derived semiochemicals. We pipetted 10 mL of the treatment stimulus [synthetic nectar broth inoculated with *L. thermotolerans, M. lactis* or *M. luteus* and incubated as described above] into a sterile 92-mm diam Petri dish which we placed into a randomly assigned delta trap. The paired control stimulus consisted of a sterile synthetic nectar broth (10 mL) presented the same way. We bioassayed the response of female *Cx. pipiens* as described above in “behavioural bioassays”.

### Attractiveness of L. thermotolerans growing on different media

In two-choice laboratory experiments with a paired-trap design, we tested attraction of female *Cx. pipiens* to *L. thermotolerans* with 40-60% surface area coverage cultured on YEPD agar, malt extract agar, or synthetic nectar agar prepared as described above. In each bioassay, the corresponding uninoculated agar media served as the paired control stimulus.

### Comparative attractiveness of single-vs multiple-species of microbes

In two-choice laboratory experiments with paired traps, we compared attraction of female *Cx. pipiens* to *L. thermotolerans, M. lactis,* and *M. luteus* presented singly or in ternary combination in the same trap. We cultured each microbe separately in synthetic nectar broth as described above, and pipetted 3.3 mL of each broth into a separate sterile 35-mm Petri dish (Sarstedt Inc., Nümbrecht, Germany). As a result, paired traps were baited with either one or three Petri dishes in each bioassay, the design of which is as described in the “Behavioural bioassay” section.

### Dose of microbes tested

We determined the concentration of microbes used in experiments by performing serial dilutions of microbe-inoculated synthetic nectar broth after incubation at 25 °C for 48 h. Cell density was determined using a hemacytometer.

### Measurement of CO_2_ production by L. thermotolerans

We inoculated sterile synthetic nectar broth (25 mL) with a single colony-forming unit of *L. thermotolerans* previously grown on YEPD agar, and then incubated the broth at 30 °C for 5 days. We added an aliquot (10 μL) of this broth to 100 mL of sterile synthetic nectar broth in a 250-mL Erlenmeyer flask, attached a vapour lock (5 mL of 98% sulfuric acid) to maintain vapour pressure and prevent water loss, and obtained the starting weight. We incubated the entire assembly in a water bath kept at 30 °C in the fume hood and monitored weight loss as a proxy for CO_2_ emission [80].

### Dynamic headspace odorant collections

We placed 12 plates with *L. thermotolerans* grown for 48-72 h at 23-26 °C on YEPD, malt agar, or SN media into a Pyrex^®^ glass chamber (34 cm high × 12.5 cm wide). A mechanical pump drew charcoal-filtered air at a flow of 1 L min^-1^ for 24-72 h through the chamber and through a glass column (6 mm outer diameter × 150 mm) containing 200 mg of Porapak-Q™ adsorbent. We desorbed odorants captured on Porapak with 0.5 mL each of pentane and ether. We analyzed 2-μl aliquots of Porapak-Q™ extract by gas chromatography-mass spectrometry (GC-MS), operating a Saturn 2000 Ion Trap GC-MS fitted with a DB-5 GC-MS column (30 m × 0.25 mm i.d.; Agilent Technologies Inc., Santa Clara, USA) in full-scan electron impact mode. To chromatograph the odorants of *L. thermotolerans* on malt extract agar and on synthetic nectar agar, we used a flow of helium (35 cm s^-1^) as the carrier gas with the following temperature program: 50 °C (5 min), 10 °C min^-1^ to 280 °C (held for 10 min). The temperature of the injector port was 250 °C and the ion trap was set to 200 °C. To analyze the odorants of *L. thermotolerans* on YEPD agar, and to reveal very volatile compounds that may have eluded detection using the above temperature program, we retained the same helium flow (35 cm s^-1^) but lowered the initial temperature, running the following temperature program: 40 °C (5 min), 10 °C min^-1^ to 280 °C (held for 10 min). Aliquots of headspace odorant extracts were injected in split mode with a 1:1 split ratio, and the temperature of the injector port and the ion trap were set to 250 °C and 200 °C, respectively. We identified odorants in headspace odorant extracts by comparing their retention indices (RI; relative to *n-*alkane standards) [81] and their mass spectra with those reported in the literature and with those of authentic standards.

### Statistical analyses

We used SAS software version 9.4 (SAS Institute Inc., Cary, USA) to analyze data, excluding from analyses all experimental replicates with no mosquitoes captured in traps. We used a binary logistic regression model with a logit link function to compare mean proportions of responders between test stimuli, and used back-transformed data to attain means and confidence intervals

## End Matter

### Ethics approval and consent to participate

None to declare

### Consent to publish

Not applicable

### Funding

The research was supported by scholarships to DP (Natural Sciences and Engineering Research Council of Canada [NSERC; https://www.canada.ca/en/science-engineering-research.html] PGSD; SFU Provost’s Prize of Distinction [https://www.sfu.ca/dean-gradstudies/awards/entrance-scholarships/provost-awards/ppd.html]; John H Borden Scholarship [http://esc-sec.ca/student/student-awards/#toggle-id-5]), scholarships to NY and EG (NSERC - Undergraduate Student Research Award), and by an NSERC - Industrial Research Chair to GG, with Scotts Canada Ltd. as the industrial sponsor.

### Author Contributions and Notes

DP, GG, MM, and CL designed the study. DP collected the microbe samples. DP, CC, HC, NY, SG, SM, AB, and EG identified the microbes. RG identified the microbial odorants. DP, EG, SM, and YU ran the bioassays. DP, EG, and YU analyzed data statistically. DP and GG wrote the manuscript. All authors reviewed and approved of the final draft.

## Acknowledgments

We thank Martin Duckhorn for his volunteer assistance in the project.

